# AZD5582 robustly reactivates latently infected cells and clears the majority of those reactivated from the SIV reservoir

**DOI:** 10.64898/2026.06.01.729198

**Authors:** Tin Phan, Maud Mavigner, Amir Dashti, Ann Chahroudi, Ruy M Ribeiro, Ruian Ke, Alan S. Perelson

## Abstract

AZD5582 (AZD) is a latency reversing agent used to support the “shock-and-kill” strategy in HIV-1 cure research. Previous studies in ART-suppressed rhesus macaques have shown that AZD can promote reactivation of latently infected cells, resulting in 2–3 log increases in on-ART viral load and significant reductions in SIV reservoir size over 5–10 doses. To quantify the impact of AZD on the reservoir, we developed an ensemble of mechanistic viral dynamic models and fit them to longitudinal plasma SIV RNA and SIV CA-DNA data from 23 macaques treated with AZD in combination with other therapies. The aggregate predictions of the model ensemble recapitulate the reactivation patterns observed in both SIV RNA and SIV CA-DNA and provide robust estimates of key parameters associated with reactivated cells. We found that AZD reactivates approximately 25% of cells in the latent reservoir per dose, with a mean reactivation duration of about 5–6 days. Of the reactivated cells, 60–79% are eventually cleared, while the remainder enter a state refractory to AZD stimulation before returning to latency. Because of this refractory state, each consecutive weekly dose reactivates about 28% fewer cells than the previous one, an effect that could be more pronounced if the refractory period substantially exceeds the interval between doses. However, the duration of this refractory state remains uncertain. Altogether, our results suggest that AZD-reactivated cells are effectively cleared. Future work should focus on improving LRAs that safely reactivate a larger fraction of the latent reservoir. Furthermore, designing experiments with varying dosing schedules can help better quantify the duration of refractoriness, which will be important for informing optimal treatment schedules and maximizing the effect of LRAs.

## I. Introduction

Despite decades of antiretroviral therapy (ART), the HIV latent reservoir persists, undermining cure efforts [1]. People with HIV, therefore, require life-long ART to control disease progression [2], posing a major barrier to its eradication worldwide [3]. If ART is interrupted, latently infected cells that spontaneously reactivate can lead to viral rebound [4–6]. The size of the latent reservoir at the time of ART interruption has been linked to the timing of viral rebound [7–10], with the magnitude of rebound likely determined by the capacity of the immune response to react to the viral reactivation [11–16]. The persistence of latently infected cells is due to their minimal HIV-1 gene expression, masking them from immune recognition and elimination [17]. Thus, the “shock-and-kill” strategy was proposed, which aims to reactivate latently infected cells with latency reversing agents (LRAs) and allow the immune response, viral cytopathic effects, and other treatments to reduce the latent reservoir [18,19]. While numerous LRAs have been evaluated for their therapeutic potential [20–24], identifying agents that robustly induce HIV transcription and meaningfully reduce the latent reservoir in vivo remains a key challenge.

AZD5582 (AZD) is a SMAC mimetic (second mitochondrial-derived activator of caspases), a potent LRA that directly inhibits the cellular inhibition of apoptosis proteins cIAP1 and cIAP2. This inhibition allows the NF-κB inducing kinase (NIK) to accumulate, leading to phosphorylation and processing of p100 into p52, formation of p52–RelB complexes, and their translocation to the nucleus [25,26]. When p52–RelB binds κB sites within the HIV-1 long terminal repeat promoter, it can activate HIV transcription and facilitate the reactivation of latently infected cells [23,27–29].

Multiple *in vitro* and *in vivo* experiments have demonstrated that AZD can robustly reactivate latently infected cells, resulting in a 2–3 log increase in on-ART viremia [30–34]. Due to its high selectivity for the non-canonical NF-κB pathway, AZD has limited off-target effects and lower immune-mediated toxicity [30,31,35] compared to drugs targeting the canonical NF-κB activation [36]. However, despite robust viral reactivation, the reduction in the size of the latent reservoir is less pronounced and appears to depend on concurrent treatments and anatomical site [32]. Several mathematical models have suggested that the relationship between viral reactivation and reservoir reduction is influenced by factors such as the characteristics of the reactivated cells and their interactions with the immune response and co-administered therapies [37–39].

Mathematical models have played an important role in understanding HIV infection and treatment responses [40–45], including those targeting the latent reservoir [37–39,46–51]. Previously, we developed models to study the impact of LRAs on the latent reservoir [37,38] and found that vorinostat, a histone deacetylase inhibitor, can increase HIV expression in latently infected cells but fails to induce substantial death of latently infected cells. Here, to study the latency reversal effect of AZD, we develop an ensemble of mechanistically linked viral dynamic models and fit them to longitudinal data from SIV-infected rhesus macaques treated with AZD in combination with other therapies. Models in the ensemble describe different mechanisms for the AZD-induced reactivation process and the interaction of reactivated cells with the immune system. By aggregating estimates across this ensemble, we aim to identify the mechanisms and quantitative characteristics of AZD-induced latency reversal to understand the impact of AZD on latently infected cells.

## II. Methods

### Data

We study the effect of AZD5582 using on-ART viremia data from three groups of adult rhesus macaques of similar age and from the same laboratory [32,33]. All macaques were infected with 3000 TCID_50_ (50% tissue culture infective dose) of SIV_mac239_ and put on ART 8 weeks post infection. After 81 – 87 weeks on ART, the first two groups of macaques received 10 weekly doses of AZD at 0.1 mg kg ^−1^ intravenously, with the interval between the 5^th^ and 6^th^ dose extended to 10 days [32]. Both groups also received four functionally distinct SIV Env-specific Rhesus monoclonal antibodies (RhmAbs) aiming to enhance viral clearance during reactivation [32]. In addition, the first group received the IL-15 superagonist N-803, an immune activator of CD8+ T cells and NK cells [52] that can potentially boost both the latency reversal effect and the elimination of infected cells [53]. Lastly, after 84 – 85 weeks on ART, the third group received a single dose of an anti-CD8α antibody (MT807R1) prior to 5 weekly doses of AZD at 0.1 mg kg ^−1^ intravenously, which may enhance latency reversal [33]. Fig. 1 summarizes the experimental designs for these groups of macaques.

**Figure 1.**
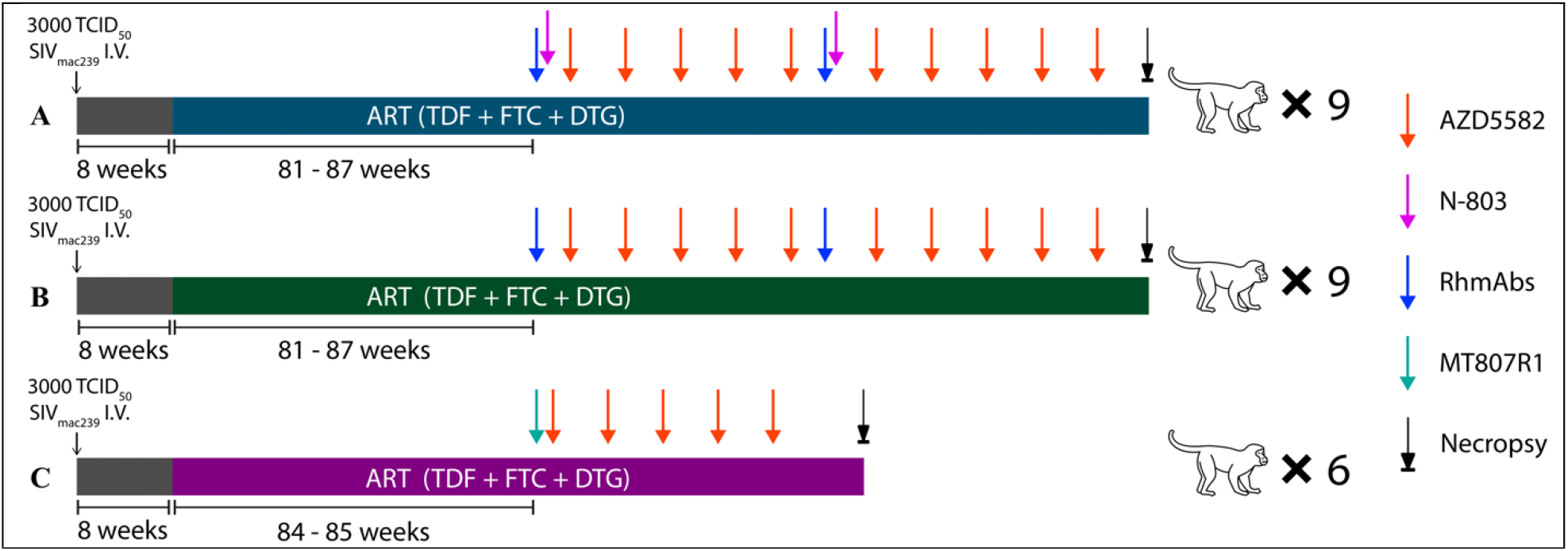
**Experimental design** from Dashti et al. [32] and Mavigner et al. [33]. All three groups of macaques were infected with 3000 TCID_50_ of SIV_mac239_ and put on ART consisting of daily dosing of 20 mg kg^-1^ subcutaneously (s.c.) tenofovir (TDF), 40 mg kg^-1^ s.c. emtricitabine (FTC), and 2.5 mg kg^-1^ s.c. dolutegravir (DTG). **(A)** After 81 – 87 weeks on ART, the 9 macaques in group 1 received weekly 0.1 mg kg^-1^ intravenously (i.v.) AZD, and twice 100 µg kg^-1^ s.c. N-803 and 20 mg kg^-1^ s.c. of each RhmAb. The N-803 and RhmAbs were given 3 days before the first and sixth AZD doses [32]. **(B)** After 81 – 87 weeks on ART, the 9 macaques in group 2 received weekly 0.1 mg kg^-1^ i.v. AZD and 20 mg kg^-1^ s.c. of each RhmAb [32]. Schedule for RhmAbs in group 2 is the same as group 1. **(C)** After 84 – 85 weeks on ART, the 6 macaques in group 3 received 50 mg kg^-1^ s.c. MT807R1 and weekly 0.1 mg kg^-1^ i.v. AZD [33]. MT807R1 was given one day before AZD. We exclude one macaque from group 2 that did not have any viral load measurement above the limit of quantification of the ultra-sensitive assay during the duration of AZD treatment to focus on the latency reversal effect when it is observable. Thus, in total, we use the data of 23 macaques for our analysis.

In addition to viral load data, we also used SIV cell-associated DNA (CA-DNA) measurements from peripheral blood to better constrain the change in the latent reservoir size following AZD. While Mavigner et al. [33] FACS-sorted CD4+ T cells to distinguish resting cells in their measurements of CA-DNA, Dashti et al. [32] used total CD4+ T cells. This difference is likely insignificant because the contribution of productively infected (non-resting) cells to total CA-DNA measurements is expected to be minimal during ART. Lastly, we used plasma AZD concentrations measured in a different set of three macaques given the same dose of AZD [30] to build a simple pharmacokinetic-pharmacodynamic (PK-PD) model for AZD.

### Mathematical Models

A major challenge in capturing the effect of an LRA using mathematical models is the high uncertainty in measurements and inferences due to the low levels of on-ART viremia in animals. Thus, there may be low confidence in a single model that best fits the small changes in the data. For this reason, we compound confidence by using an ensemble of mechanistically linked models, where the models share a core structure but differ slightly in the reactivation process, or the interaction of the reactivated cells with the immune response. Ensemble modeling is a powerful strategy to provide more robust estimates and predictions in the presence of unknowns and uncertainty. It is frequently used in epidemiology [54] and weather forecasting [55]. Existing applications of ensemble modeling focus on model prediction. Previous studies comparing multiple related viral dynamic models often show similar estimates for shared parameters [11,56–60], which underlies the possibility of estimating shared parameters accurately using an ensemble of mechanistically linked models, and thus provide more robust conclusions about the underlying biological mechanisms.

Here, we considered a class of within-host virus-immune models resulting from the modeling framework introduced by Conway and Perelson [9], which has been shown to capture HIV and SIV viral load trajectories under different scenarios [11,61–63]. In the base model, the dynamics of latently infected cells, *L*, is governed by their per capita death rate *d*_*L*_, activation rate *a*, and proliferation rate *ρ*. The net effect of these processes results in a half-life *t*_1/2_ for latently infected cells of approximately 44 months [1,64,65]. Productively infected cells, *I*, produce virus at rate *p* per cell and die at per capita rate *δ*_*I*_. Virus is cleared at per capita rate *c*. Effector cells, *E*, are produced at rate *λ*_*E*_, die at per capita rate *d*_*E*_, and expand upon contact with productively infected cells at rate 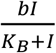 [9,11,66,67], where *K*_*B*_ is the number of infected cells required for the effector cell population to expand at half its maximal rate *b*. Effector cells kill productively infected cells [68,69] at rate *m*_1_*I* and reduce viral production, via noncytolytic means [70–74], by a factor 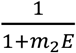. The inclusion of an effector cell compartment is based on previous studies suggesting that CD8+ T cells play a central role in the control of HIV/SIV even during ART. For example, an experiment involving 13 SIV-infected macaques showed that while the macaques were on ART, the depletion of CD8+ T cells resulted in rapid viral rebounds, which were controlled once CD8+ T cells repopulate [62,75]. The equations describing the base model during ART are

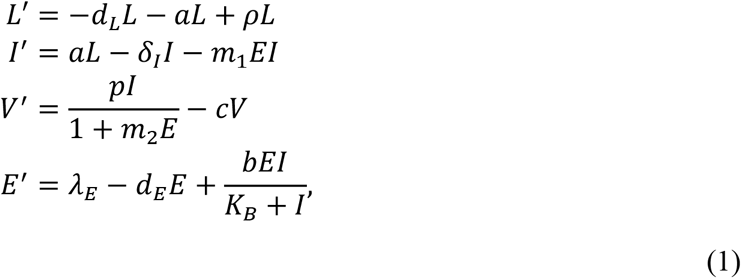

where we have assumed that ART is sufficiently effective that new infections can be ignored and hence an equation for target cells is not needed.

We model the plasma concentration of AZD, *Z*(*t*), using a one-compartment pharmacokinetics (PK) model with infusion rate constant *k*_*a*_ over a duration Δ_*t*_ and tissue distribution rate constant *k*. The function *D*(*t*) denotes the administration schedule of AZD.

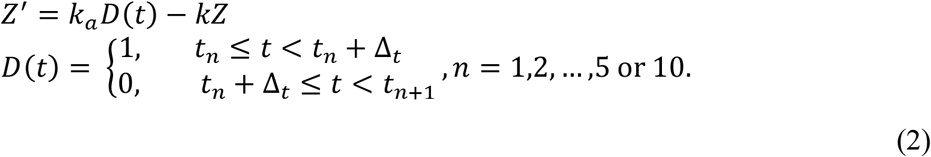

We estimated the PK parameters by fitting *Z*(*t*) to the longitudinal plasma concentrations of AZD following a single infusion in three macaques [30], see details in S1 text and Fig. S1. To integrate the PK model within the larger framework, we fixed the absorption and elimination rate constants *k*_*a*_ and *k* to the medians of the three individual estimates, namely 5 × 10^4^ nM/day and 90/day, respectively and Δ_*t*_ is 1/48 day (Table S1). With these fixed PK parameters and using the molecular weight conversion of 1015.29 g/mol for AZD [76], each macaque reaches a maximum concentration of ~465 nM, which is within the range measured [364 – 630] ng/mL [30]. We model the pharmacodynamics (PD) of AZD using an E_max_ model 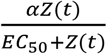 [77], where *EC*_50_ is the half-maximal effective concentration, taken to be 7.5 nM [31], and *α* is the maximal latently infected cell reactivation rate due to AZD (and can be different from the natural reactivation rate, *a*). By fixing the PK and PD parameters, we assume that differences in reactivation patterns among the treated macaques are not primarily driven by differences in the individual sensitivity to the drug’s effect.

In the first model variation (Fig. 2A), AZD directly activates latently infected cells into productively infected cells at per capita rate 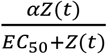. Thus, the equations for *L* and *I* are changed to

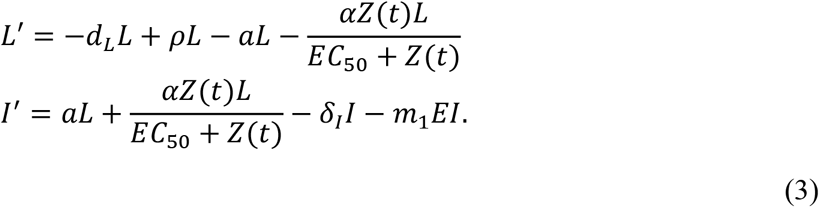

**Figure 2.**
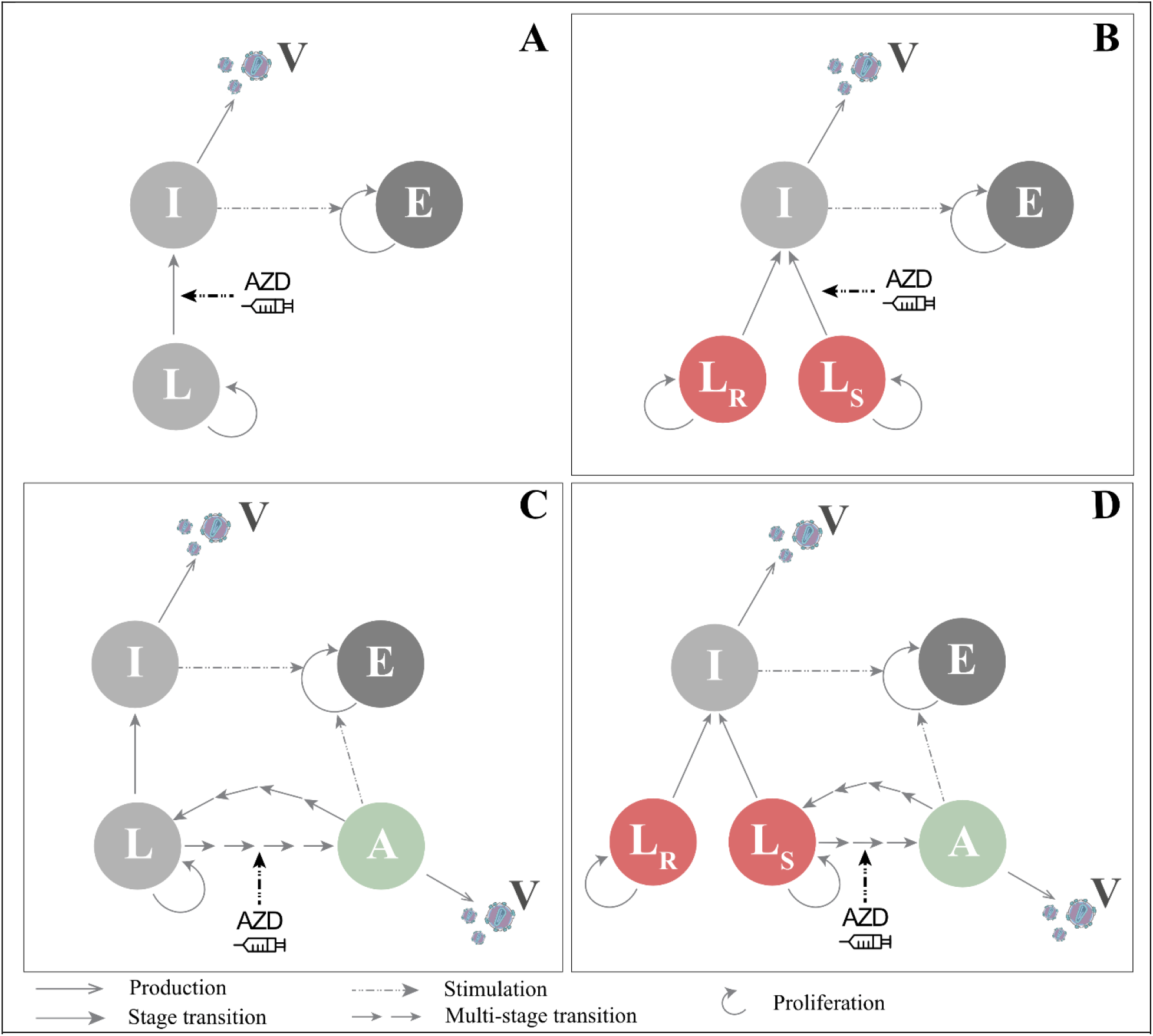
Main model variations. V: viruses; L: latently infected cells; I: productively infected cells; A: AZD-reactivated cells; E: Effector cells. **(A)** The first variation assumes AZD directly activates latently infected cells into productively infected cells. **(B)** The second variation assumes AZD only reactivates a subset of latently infected cells (L_S_) into productively infected cells, while the remaining latently infected cells (L_R_) are resistant to its effect. **(C)** The third variation assumes AZD-reactivated cells differ from productively infected cells. AZD-reactivated cells may also enter a refractory state before returning to latency. **(D)** The fourth variation combines the second and third variations.

In the second variation (Fig. 2B), we consider the possibility that a population of latently infected cells are intrinsically resistant to AZD activation, which could be due to integration of the virus into a region of the genome not responsive to non-canonical NF-κB activation [26,78– 80], or deep silencing by repressive histone marks [81,82]. Hence, we model two distinct categories of latently infected cells, one susceptible, *L*_*S*_, and the other resistant, *L*_*R*_, to the effects of AZD. The equations that are changed for this model variation (Fig. 2B) are

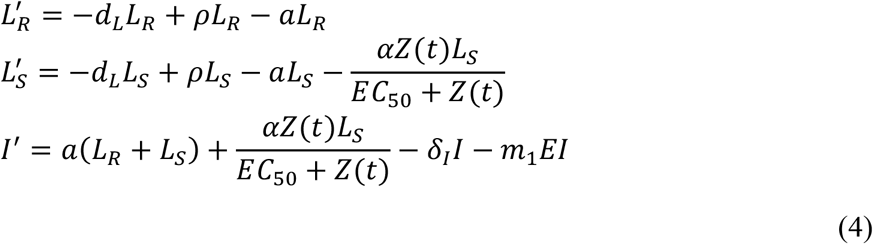

In the two models above, we assume that after reactivation by AZD, latently infected cells (*L*) become productively infected cells (*I*). In the third variation (Fig. 2C), we assume that cells reactivated by AZD (*A*) behave differently from productively infected cells (*I*), as we proposed before [37,38]. More specifically, AZD-reactivated cells are assumed to produce virus, induce effector cell expansion, and be susceptible to immune killing and cytotoxic effects at a fraction *ϵ*_*A*_, termed the relative reactivation efficiency ratio, of the rates of productively infected cells [38,83]. The extension that incorporates *A* is

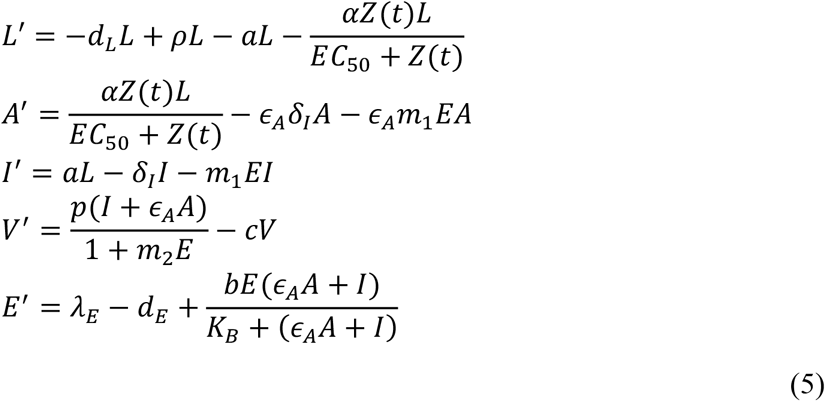

An additional consideration is that AZD-reactivated cells may return to latency [84] or enter a refractory state – not susceptible to further induction by AZD5582 – before returning to latency. We previously found that this refractory state was needed to explain viral load data under vorinostat [38]. The refractory state could account for factors affecting the cell susceptibility to AZD after the initial activation. For example, the time required to reset pathway sensitivity following sustained drug-induced reactivation, as the non-canonical NF-κB pathway involves numerous tightly regulated molecules [26], with key components such as NIK being depleted in the process [85–87]. For similar reasons, non-canonical reactivation likely occurs slower than canonical reactivation [36]. Thus, we model reactivation and returning-to-susceptibility as state transitions with gamma-distributed waiting times [38,88]. Upon AZD-induced reactivation, latently infected cells *L*_0_ go through *N* intermediate reactivation states *L*_*i*_ (*i* = 1, …, *N*) at rate *γ* before becoming fully reactivated cells, *A*, that produce virus. The fully reactivated cells return to latency at rate *ω*, going through *M* intermediate states *R*_*j*_ (*j* = 1, …, *M*) that are refractory to further drug effect, prior to returning to the drug-susceptible latent state *L*_0_. Together, the model equations for this variation are (Fig. 2C)

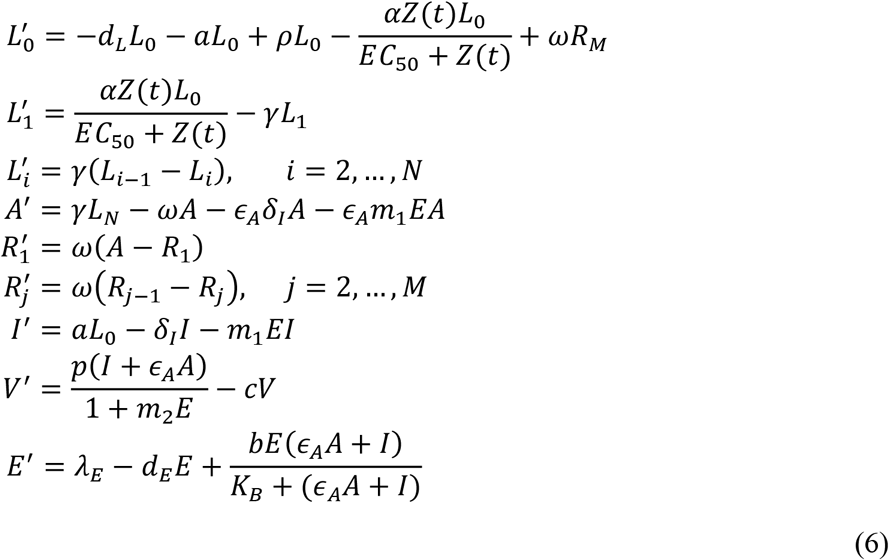

The last model variation (Fig. 2D) combines the second and third variations. Thus, it includes AZD-resistant and -sensitive latent cells, stepwise reactivation, and a refractory state.

We model the effect of the anti-CD8α monoclonal antibody MT807R1 as a constant 10-fold increase in the death rate of effector cells once the drug is administered, since CD8+ T cells were depleted for almost the entire duration of AZD treatment [33]. The magnitude of the increased death rate is roughly consistent with that found by Cao et al. [62] for CD8 depletion with the same antibody in a different experiment.

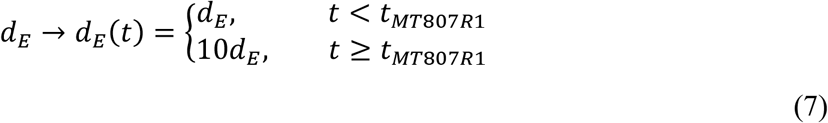

In addition to the four main model variations described above, we examined in the supplementary material whether the model fit can be improved by including viral infection, varying the number of activation and refractory stages, fixing or removing random effects on certain parameters such as *L*_0_(0), or specifying in more detail how AZD-reactivated cells might differ from productively infected cells, e.g., separating *ϵ*_*A*_ into multiple terms 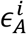 for different processes (viral production, cell death, immune stimulation, etc.). For instance, to test the possibility of a small amount of de novo infection as a result of latency reversal (although unlikely with ART), we add a source of production of latently and productively infected cells at rates *β*_*V*_*V, f*_*L*_β_*V*_*V* and (1 − *f*_*L*_)β_*V*_*V*, respectively, where *β*_*V*_ is the infection rate assuming the number of target cells remains relatively constant on ART and *f*_*L*_ is the fraction of infections that generate latently infected cells (see M1 in S1 Text). In this version, we also examine the possibility that effector cells may release cytokines or chemokines to reduce the infection rate by 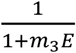. Thus, we tested families of alternative models based on each of the variation shown in Figure 2.

### Model fitting and comparison

We used a nonlinear mixed effect modeling approach (software Monolix 2024, Lixoft, SA, Antony, France) to fit the predicted viral load, log_10_ *V*, of each model to the log_10_ SIV plasma viral load data. We fit SIV CA-DNA data concurrently using 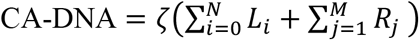, where *ζ* is a scaling constant (e.g., some CA-DNA+ cells may carry defective virus and not be replication-competent latently infected cells) subject to random effects to account for differences among the animals, which can affect CA-DNA measurements. In model variations that separate AZD-susceptible (*L*_*S*_) and AZD-resistant (*L*_*R*_) population, we use 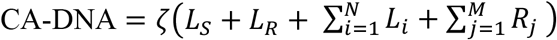. Here, we assume the number of virus-producing cells, *I* and *A*, is small compared to the number of latently infected cells, *L* and *R*, during ART, which is shown to be consistent with the prediction of the best model, hence can be neglected when calculating CA-DNA. We also examined 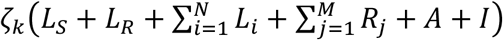 and 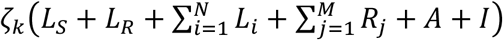, which gave similar results. The scaling factor *ζ*_*k*_ is a constant for each treatment group (*k* = 1,2,3) due to the assumption that the fraction of replication-competent latently infected cells susceptible to reactivation by AZD is relatively stable for the 5–10-week treatment duration. We applied left censoring for values at the limit of quantification. Fixed parameters and the ranges for all fitting parameters are provided in Table S2. All fitting parameters have random effects. Initially, we choose the initial guesses for the parameters manually to avoid unrealistic model dynamics. For consistency, all final results use the same initial guesses.

We assume the system is at steady state prior to AZD treatment and approximate the initial conditions under the assumption that the expansion of effector cells induced by productively infected cells is negligible during ART and prior to AZD treatment [11]. A summary of the description, range, and reference for all parameters is provided in Table S2. Model comparison was done using the corrected Bayesian Information Criterion (BICc) [89] as reported by Monolix.

## III. Results

### The model ensemble fits the data well and generates uniform parameter estimates

We developed and studied 31 models based on the four model variations described in the Methods. The description of the 31 model variations is given in S2 Text. Table S3 and Fig. 3A quantify the model fitting results, including the negative log likelihood (−2LL) and BICc, for all 31 models. The model with the lowest negative log likelihood will have the best fit, but to prevent overfitting, we penalize models with additional parameters using the BICc. The preferred model is M26, the model with the lowest BICc. It belongs to the family of models shown in Fig. 2C (the third main model variation, Equation 6), and its fit to the data is shown in Fig. 4A. To further examine the key features of AZD reactivation, we selected 10 representative models (indicated by the red asterisks in Fig. 3A), chosen to have low values of BICc with distinct features (i.e., mechanisms) from one another. For example, M23-27 all have low BICc, but they are mechanistically the same, except for the number of transitioning stages (L ↔ A), so we select only the best of these models (M26) as a representative of this group of models. Hereafter, we refer to these 10 models collectively as the *model ensemble*. The distributions of parameter estimates from the model ensemble are shown in Fig. 3B, showing population estimates with the [Q1, Q3] range falling within one order of magnitude or less. The corresponding values are listed in Table S4.

**Figure 3.**
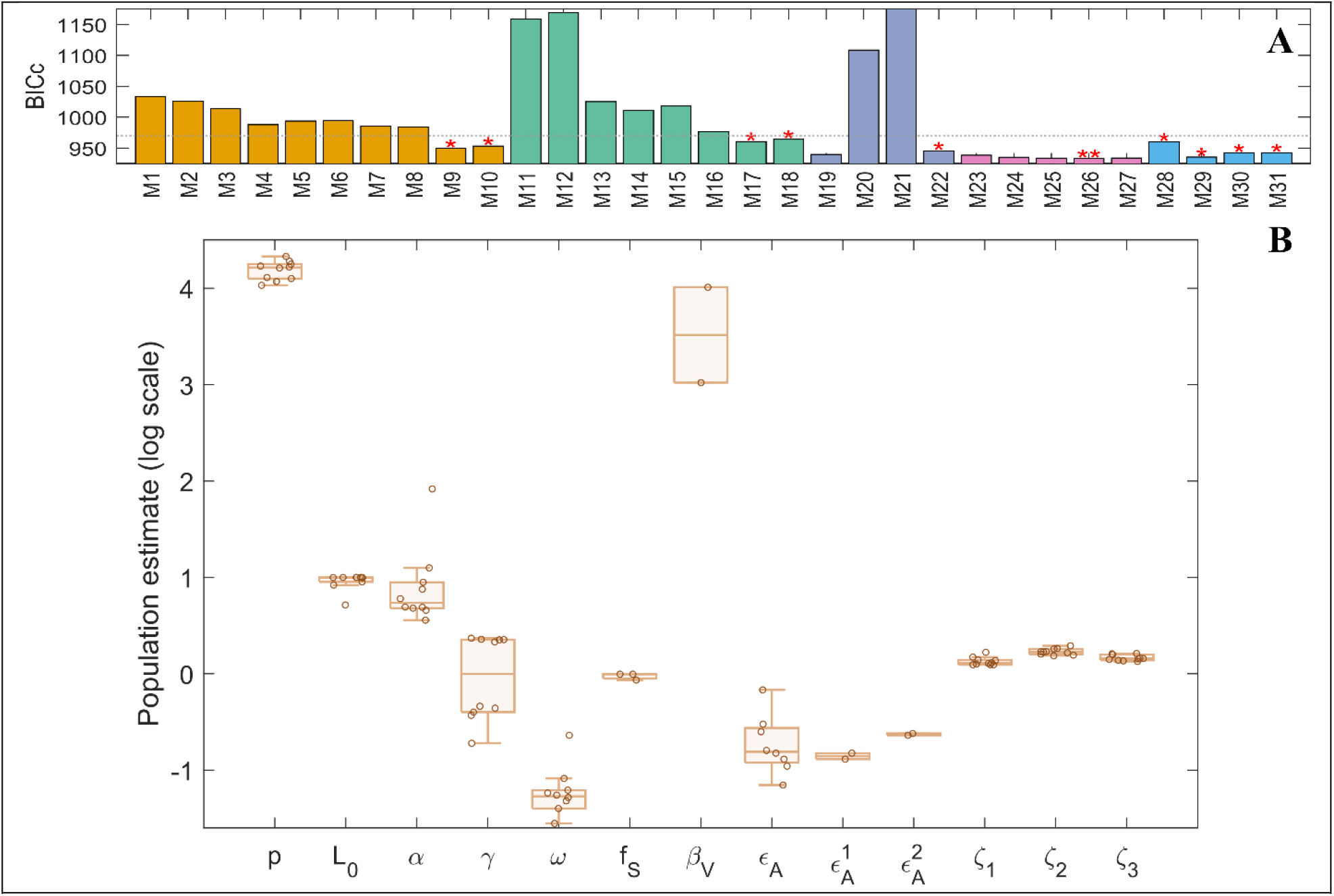
Model ensemble summary. (**A**) BICc of 31 model variations (M1 to M31). Orange bars: main model variations with or without effector cells. Teal bars: tests for the inclusion of active infection. Blue bars: test of M9 with either *L*_0_(0) fixed at different values or without random effect. Pink bars: test of M22 with varying transition stage numbers N and M. Cerulean bars: test for robustness of M26. The red asterisks denote the selected models to be included in the ensemble analysis. Horizontal dotted line shows a cut-off BICc of ~968 (or about 35 ΔBICc) for the selection of the ensemble models. Two asterisks on M26 denote the best model by BICc. (**B**) Distribution of population parameter estimates for the model ensemble. The lower and upper limits of the boxplot represent the first and third quartiles, respectively. The line inside the box is the median, and the whiskers – when present - connect the top/bottom of the box to the max/min values that are not outliers (data points further than 1.5 times the interquartile range). Overlaid circles are individual model estimates. p is the viral production rate, L_0_ is the initial size of latent reservoir, α is the maximum reactivation rate of the noncanonical NF-κB pathway by AZD, γ is the activation transition rate, ω is refractory transition rate, f_S_ is the fraction of the latent reservoir susceptible to activation by AZD, β_V_ is the infection rate constant, ϵ_A_ is the relative reactivation efficiency ratio, 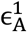 is the relative reactivation efficiency ratio specific to cytotoxicity, 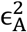 is the relative reactivation efficiency ratio specific to viral production, ζ_k_ (k = 1,2,3) are the group-specific constant scaling to CA-DNA.

**Figure 4.**
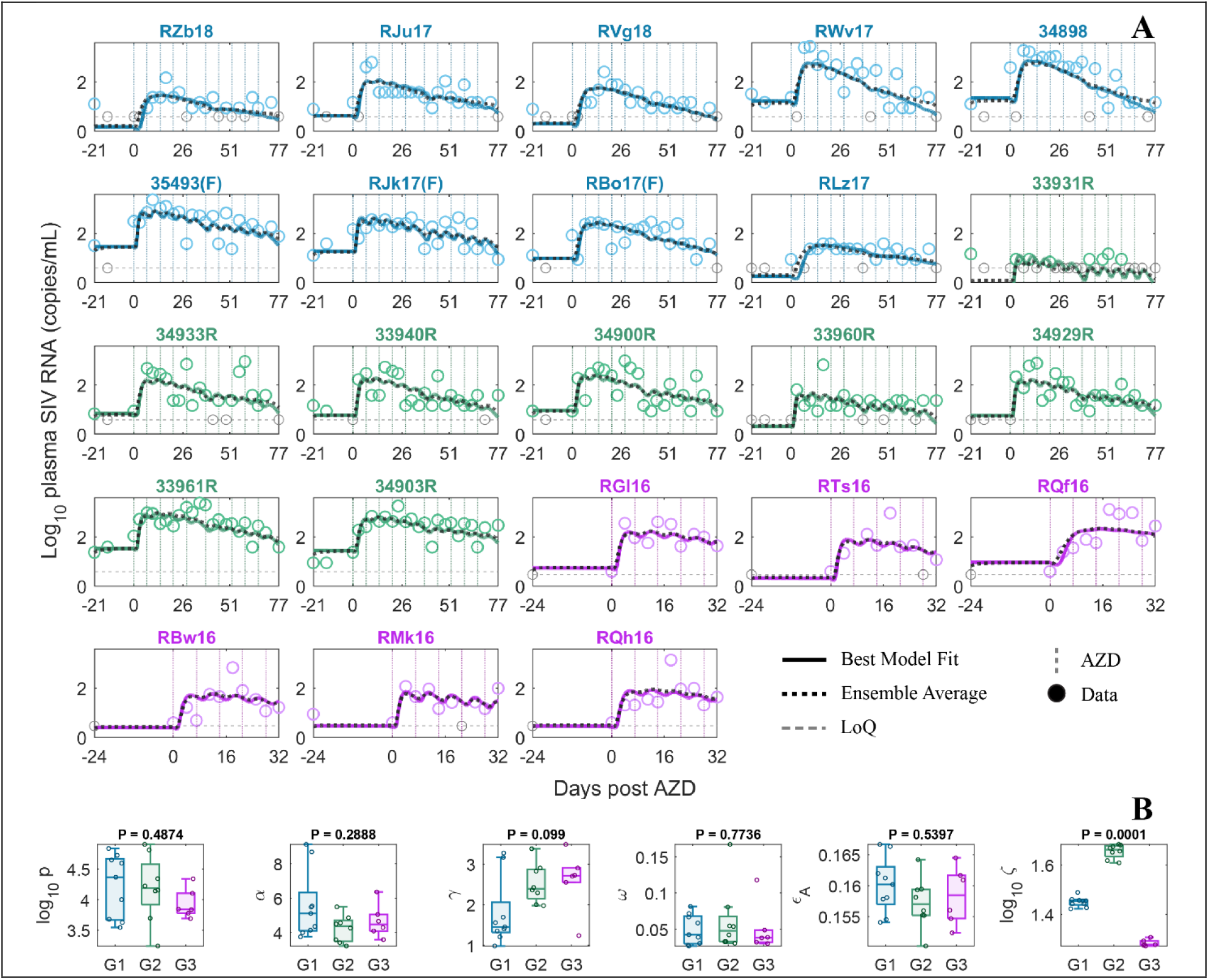
Best model fit and group parameter estimates. (**A**) Best model (M26) fit to viral load data (solid curve). Blue is the group treated with AZD, N-803, and RhmAbs. Green is the group treated with AZD and RhmAbs. Purple is the group treated with anti-CD8α prior to AZD. Ensemble average is represented by the black dashed curve. Gray circles are data points below the limit of quantification (dotted horizontal gray line). Non-gray circles are data points above the limit of quantification. The dotted vertical lines indicate AZD administrations. (**B**) Summary of best fit parameters for M26. P-values are obtained from the Kruskal-Wallis test. G1 is the group treated with AZD, N-803, and RhmAbs. G2 is the group treated with AZD and RhmAbs. G3 is the group treated with anti-CD8α prior to AZD.

### AZD-reactivated cells likely differ from productively infected cells

Collectively, models that assume AZD-reactivated cells differ from productively infected cells (3^rd^ and 4^th^ model variations) have the lowest BICc. Interestingly, models of the 4^th^ variation, which is just the 3^rd^ model variation with the added assumption that some latently infected cells are not susceptible to AZD, always fit the viral load data better than their 3^rd^ variation equivalent (i.e., lower −2LL), but have a worse BICc due to additional parameters — as seen in M5 vs. M4, M10 vs. M9, M15 vs. M14, M18 vs. M17, M30 vs. M26, and M31 vs. M29 (Table S3). On the other hand, models that only assume the existence of an AZD-resistant subpopulation of latently infected cells without distinguishing AZD-reactivated cells, *A*, from productively infected cells, *I*, (the 2^nd^ main variation) do not fit the data as well and have a higher BICc. Models with neither assumption (the 1^st^ main variation) fit the data the worst. These results support a distinction between AZD-reactivated cells and productively infected cells, and the possible existence of an AZD-resistant subpopulation of latently infected cells.

The inclusion of an effector cell population in the model slightly improves the model fit but substantially worsens its BICc score. Hence, none of the models with an explicit compartment for effector cells were included in the top 10 models, and they all had BICc difference (ΔBICc) larger than 50 points in relation to the best model (Table S3). This result suggests that the dataset does not allow us to explicitly estimate the impact of effector cells on the AZD reactivated cells. Furthermore, we fitted models M4 and M5 and their counterparts without effector cells (M9 and M10) to data from only the two groups of macaques treated with RhmAbs. In both scenarios, the inclusion of effector cells did not improve the model fit and significantly worsened the BICc. Altogether, this suggests the inclusion of an explicit effector cell compartment in our models is unnecessary to describe this data set.

The model ensemble estimates the relative reactivation efficiency ratio *ϵ*_*A*_ to be 0.16 with [Q1, Q3] = [0.13, 0.26] (Fig. 3B), implying that on average AZD-reactivated cells produce viruses, die by cytotoxicity, and interact with the immune system at about 16% of the corresponding rates for productively infected cells. Note that *ϵ*_*A*_ represents a single summary comparison between reactivated and productively infected cells. When separately considering the relative death rate and viral production rate between AZD-reactivated cells, *A*, and productively infected cells, *I*, using 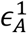 and 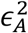, respectively (M29 and M31), 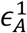 is estimated to be 0.13 – 0.15 and 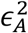 is estimated to be 0.23 – 0.24. Another interesting observation is that the median half-life for AZD-reactivated cells given by log(2) /(*ϵ*_*A*_*δ*) is 4.5 days with [Q1, Q3] = [2.6, 5.5] days. This estimate is similar to the half-life of cells with intact SIV proviruses of 3.3 days (95% CI: 2.5 – 4.5 days) within the first 4 weeks of ART found in another study [90] (consistent with the observation that most CA-DNA measured in these macaques is intact [32,33]), and the half-life of long-lived SIV productively infected cells, which was estimated at 2.5 – 6.2 days [71]. For the model ensemble, if we only consider that reactivated cells either die or return to latency, then we can approximate that the fraction of AZD-reactivated cells eventually killed is 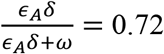 (median) with [Q1, Q3] = [0.60, 0.79].

### AZD-reactivated cells may enter a state refractory to drug induction

The inclusion of multi-stage reactivation and a refractory phase post AZD reactivation universally improves model fits, consistent with findings from our previous work on vorinostat [38]. For instance, models with both multistage reactivation and a refractory phase (M4 and M9) directly outperform their counterparts with single step reactivation (M3 and M8) by 25 and 34 BICc points, respectively. Moreover, when we remove from the best model (M26) the refractory phase while still allowing AZD-reactivated cells to return to latency (immediately), the model performance drops by over 26 BICc points (M29 vs. M26). This suggests that AZD reactivates latently infected cells in a multistage process, and supports the notion that, after reactivation, these cells enter a temporary state that is refractory to further induction by AZD.

The model ensemble robustly estimates the AZD-induced maximal reactivation rate *α* (median: 5.47, [Q1, Q3] = [4.82, 8.55] per day), the reactivation transition rate between reactivation stages *γ* (median: 1.30, [Q1, Q3] = [0.41, 2.26] per day) in models with multistate reactivation, and the refractory transition rate *ω* (median: 0.054, [Q1, Q3] = [0.042, 0.061] per day). This corroborates the capacity of AZD to activate the non-canonical NF-κB pathway and drive latency reactivation, in alignment with the substantial evidence for its efficacy both *in vitro* and *in vivo* [30–33]. The expected reactivation time, or the time it takes for a cell to be fully reactivated after initiation, is *N*/*γ*, where *N* is the number of intermediate reactivation stages. Similarly, the expected refractory duration is *M*/*ω*, where *M* is the number of intermediate refractory stages. The median expected reactivation time for the model ensemble is 5.6 days with [Q1, Q3] = [4.4, 7.3] days. This is consistent with the observed delay between the first AZD dose and viral recrudescence (Fig. 4A) and in line with the expectation that the non-canonical activation of the NF-κB pathway is slower than that of the canonical pathway [36], with canonically activated cells *in vitro* showing sign of activation within hours post stimulation [91,92]. Interestingly, this is also substantially slower than activation via histone deacetylase inhibitors [18,24]. The expected duration that AZD-reactivated cells produce viruses is given by 1/(*ω* + *ϵ*_*A*_*δ*_*I*_), which has a median of 5.0 days with [Q1, Q3] = [4.1, 5.4] days. Without considering cell death, these reactivated cells are expected to stay active for a duration of 1/*ω* before going into a refractory state, which is estimated to be 18.7 days (median) with [Q1, Q3] = [15.3, 23.8] days. This is similar to the estimated time to return to rest for activated T cells *in vitro* of 7 – 21 days [84]. Note that this is a comparable estimate because when a reactivated cell enters the first refractory state (*R*_1_), it effectively returns to a latent state as it no longer produces virus. The median expected refractory duration for the model ensemble, *M*/*ω*, is 115 days ([Q1, Q3] = [55, 197] days), suggesting a prolonged refractory period. However, this is likely an overestimate, as the weekly dosing intervals in our data are far shorter than this estimated duration, meaning subsequent doses are administered while many cells are still refractory, leaving the data largely uninformative about when refractoriness actually resolves.

### Best model

Model M26 recapitulates the observed data well (Fig. 4A) and has the lowest BICc (Table S3). It is part of the third variation (Fig. 2D) with N=M=10 and without an explicit effector cell compartment.

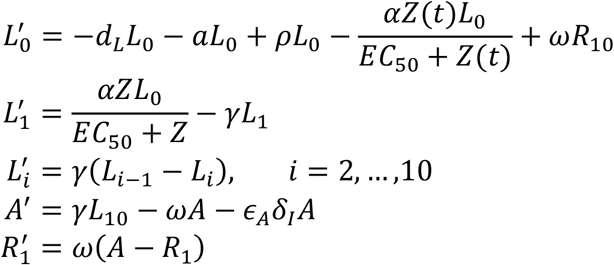

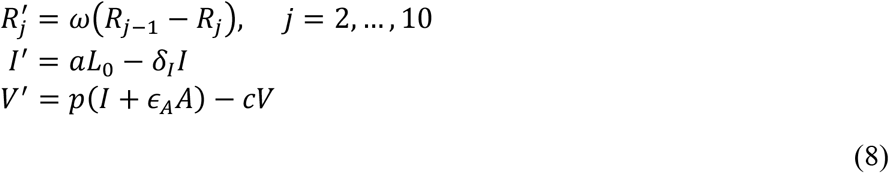

Parameter estimates obtained from the best fit model are consistent with the model ensemble estimates. Specifically, parameter population estimates of this best fitting model fall within 1.5 times of the interquartile range (or the whisker range, Fig. 3B) of the ensemble estimates and the best model fit is very similar to the fit of the model ensemble average (Fig. 4A). A spaghetti plot showing the fit of each individual model in the ensemble model to the data is shown in Fig. S3). The consistency between the best model and the ensemble reinforces the reliability of parameter estimates. Comparison of parameter distributions among treatment groups using the Kruskal-Wallis test shows no significant difference in parameter distributions among the three treatment groups, except for the individual CA-DNA scaling parameter *ζ*. However, the difference is small, i.e. 0.2 – 0.4 log (Fig. 4B).

### The impact of AZD on the latent reservoir

A refractory state post reactivation would imply that consecutive weekly AZD dosing is not optimally efficient [37]. We use model M26 and best-fit parameters for each macaque to examine the efficacy of AZD. Fig. 5A shows M26 fits to CA-DNA. Over the dosing interval, the peripheral blood CA-DNA level on average decreases by 0.61 log for the two groups receiving 10 doses and 0.47 log for the group receiving 5 doses. Note that without AZD, CA-DNA levels stay approximately constant over this duration due to the 44-month reservoir half-life. We also estimate the total number of cells reactivated for one week following each consecutive dose of AZD (Fig. 5B). A discrete exponential decay function, 3.41*e*^−0.33*n*^ with *n* denoting the n^th^ consecutive weekly dose, fits the median cumulative reactivations well (Fig. 5B). This shows that each consecutive weekly dose reactivates roughly 28% fewer cells relative to the previous dose. We also simulate the latent reservoir dynamics (Fig. 5C), which mirrors the dynamics and reduction observed for CA-DNA.

**Figure 5.**
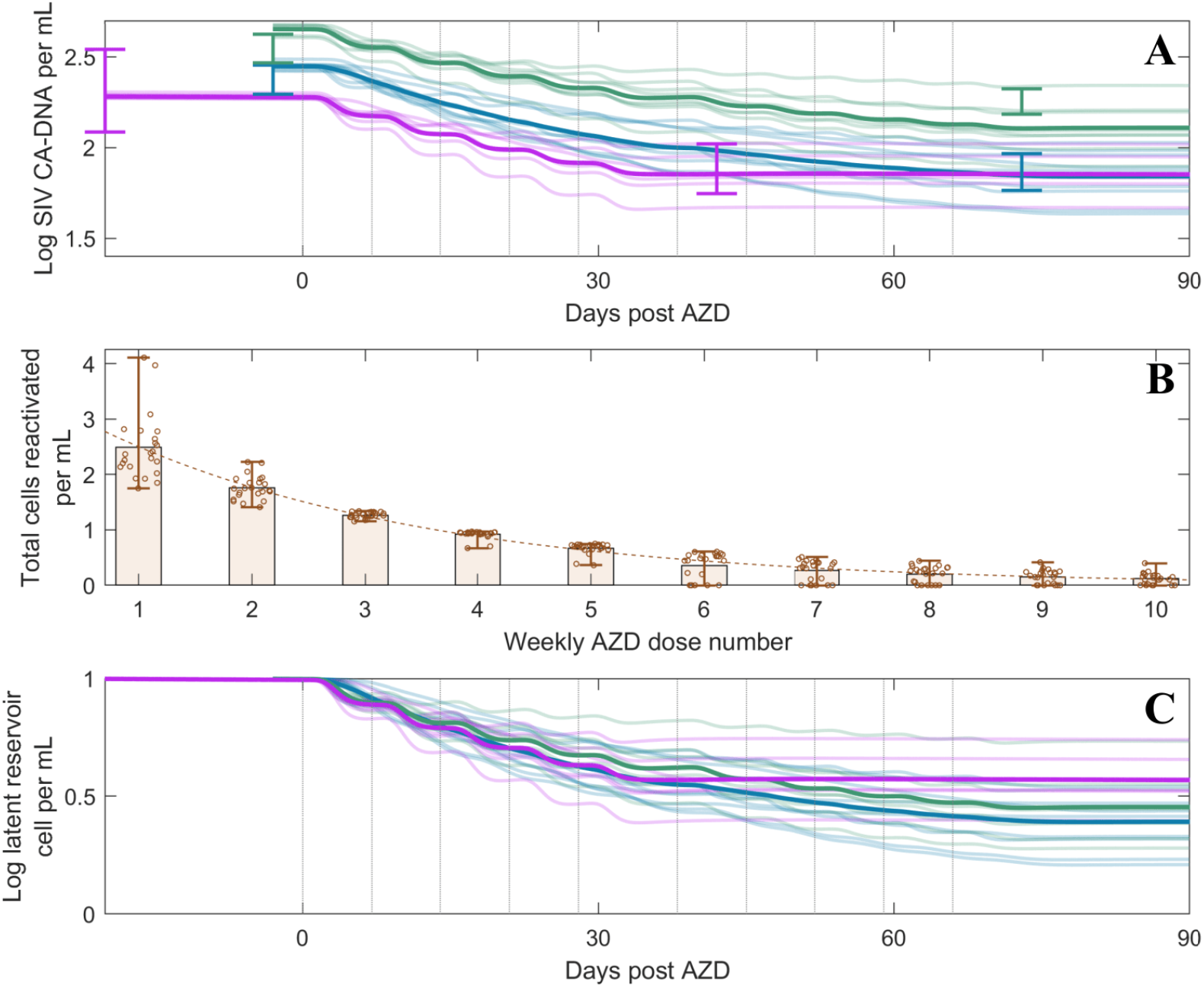
Latency reversing effect of AZD. Blue is the group treated with AZD, N-803, and RhmAbs. Green is the group treated with AZD and RhmAbs. Purple is the group treated with anti-CD8α prior to AZD. Bolded lines show population means, while thin lines show individual trajectories. **(A)** Dynamics of SIV CA-DNA, e.g., 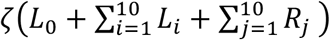. Error bars show measurement means and standard errors. The dotted vertical lines indicate AZD administrations. **(B)** Estimates for the total number of latently infected cells reactivated for seven days following each consecutive weekly dose of AZD. The lower and upper limits of the error bars show the minimum and maximum of the individual estimates (overlaid circles), respectively. For the latter doses 6^th^ to 10^th^, these estimates only include the first two groups (since the third group only had 5 doses). Dashed curve shows an exponential decay function fit to the median values. **(C)** Corresponding dynamics of the latent reservoir.

## IV. Discussion

Despite robust reactivation of latently infected cells, AZD treatment resulted in modest (~0.5–0.6 log) reductions of the latent reservoir over the 5- or 10-week dosing interval [32]. To estimate the impact of AZD on the latent reservoir, we combined longitudinal data from several macaque experiments with an ensemble of mechanistically linked viral dynamic models. We fit all models to plasma viral load and SIV CA-DNA during AZD treatment and aggregated parameter estimates from 10 best-performing models. While parameters unique to individual models varied, key parameters associated with AZD-induced latency reversal were consistent across the ensemble. The models consistently estimate that AZD reactivates a fraction (approximately 25%) of latently infected cells. Among the reactivated cells, approximately 72% eventually die while the remaining cells enter a temporary state refractory to AZD before ultimately returning to latency. Together, these results indicate that AZD5582 robustly reactivates latently infected cells and clears the majority of those reactivated. However, the existence of a refractory state suggests individuals may benefit from a longer interval between consecutive doses, although the optimal dosing interval cannot be estimated with current data.

Latently infected cells, while effectively reactivated by AZD, are intrinsically different from productively infected cells. [26,78–82]. This is likely because latently infected cells are resting cells, and drug-induced reactivation may generate a cell state that is less transcriptionally active than that of productively infected cells. Lower transcription means fewer virions produced and therefore reduced viral cytopathic effects. At modest transcription levels, proteins such as Nef, Tat, and Vpr can be anti-apoptotic, whereas pro-apoptotic effects emerge only at higher expression levels [93], so maintaining viral gene expression below this threshold may allow AZD-reactivated cells to survive longer. Reduced viral transcription is also expected to lead to lower antigen presentation at the cell surface, making these cells less likely to stimulate a strong immune response or to be efficiently targeted by immune-mediated killing. This weak interaction between LRA-reactivated cells and the immune response may help explain the similar immune-related parameter estimates across the three macaque groups, despite their exposure to different immune modulators (Fig. 3B). Notably, a recent study combining AZD with an mRNA SIV-gag vaccine in ART-suppressed macaques found that AZD may diminish effector activity of CD8+ T cell responses [94], which may be another factor contributing to the weak effector cell response estimated in our models. That said, Dashti et al. noted an increase in the SIV IFN-γ response in AZD + RhmAbs treated macaques, suggesting the immune response generated by AZD may vary by treatment [32]. We also found that explicitly including in our models de novo infection by virus produced from AZD-reactivated cells, which could in principle generate more productively infected cells and thereby stimulate stronger immune responses, did not improve the model fit. This may be because ART blocks most new infections, evidenced by the lack of long-term viral evolution in these animals [95].

The model ensemble suggests that non-canonically reactivated cells, as they return to latency, likely enter a temporary refractory state that is not responsive to further drug-induced activation. A refractory state following activation has also been proposed for another LRA, vorinostat [37,38]. For AZD, such refractoriness is likely related to the highly regulated nature of non-canonical NF-κB activation [26] and the time required to replenish molecules essential for this pathway [85–87]. Ensemble estimates place the duration of refractoriness on the order of weeks to months, implying that the fraction of latently infected cells that are refractory to AZD can progressively build up over a weekly dosing schedule. To illustrate its implication, suppose the duration of refractoriness is much longer than the dosing interval and that each dose of AZD reactivates a fraction *x* of the latently infected cell population. Then the proportion of latent cells remaining after *n* doses administered within a single refractory period is (1 − *x*)^*n*^. In this regime, rapid dosing inevitably yields lower efficiency: as fewer total number of cells are reactivated with each consecutive treatment, and higher effectiveness (larger *x*) further accentuates this diminishing-return effect. Although AZD is generally associated with low toxicity [30,31,35], prolonged or more intense exposure could still increase toxicity risk. If instead dosing is spaced relative to the duration of refractoriness, the overall effectiveness of AZD could be preserved while maintaining a low toxicity profile. Our results suggest that a compound-interest strategy with an appropriate waiting period is feasible for AZD, especially if the refractory duration can be minimized. However, a potential obstacle to this strategy is the possibility of a subset of latently infected cells that are not susceptible to the drug, or of sanctuary sites for latently infected cells, possibilities that we cannot entirely rule out, even though models with cells resistant to reactivation performed worse.

Our study has several limitations. First, the ensemble fit did not capture the clear divergence in the viral load trends in some macaques such as the high, out-of-trend second viral peaks in macaques 34933R and 33961R. This is in part due to the use of a population fitting approach, where the population trend is prioritized over individual trajectories. We suspect these high viral peaks are caused by the reactivation of a large number of latently infected cells that can generate a large quantity of virus. However, these events are likely highly stochastic and are difficult to capture with a deterministic model or even with a stochastic model. Second, the ensemble is representative of key mechanisms but not comprehensive. Nonetheless, the main features we infer are shared by all top-performing models, suggesting our main conclusions are not model-specific. Fig. 6 synthesizes key model developments and rationales, showing why the main mechanisms (possibility 4) are necessary to explain experimental observation with AZD. Future studies with sufficient data to estimate the time cells spend in the refractory state could examine whether increasing AZD or SMAC mimetic dosage or dosing frequency, and combining them with complementary drug types, can induce a more durable, robustly reactivated state that allows these cells to be eliminated by virus- and immune-mediated cytotoxic effects and antiviral drugs.

**Figure 6.**
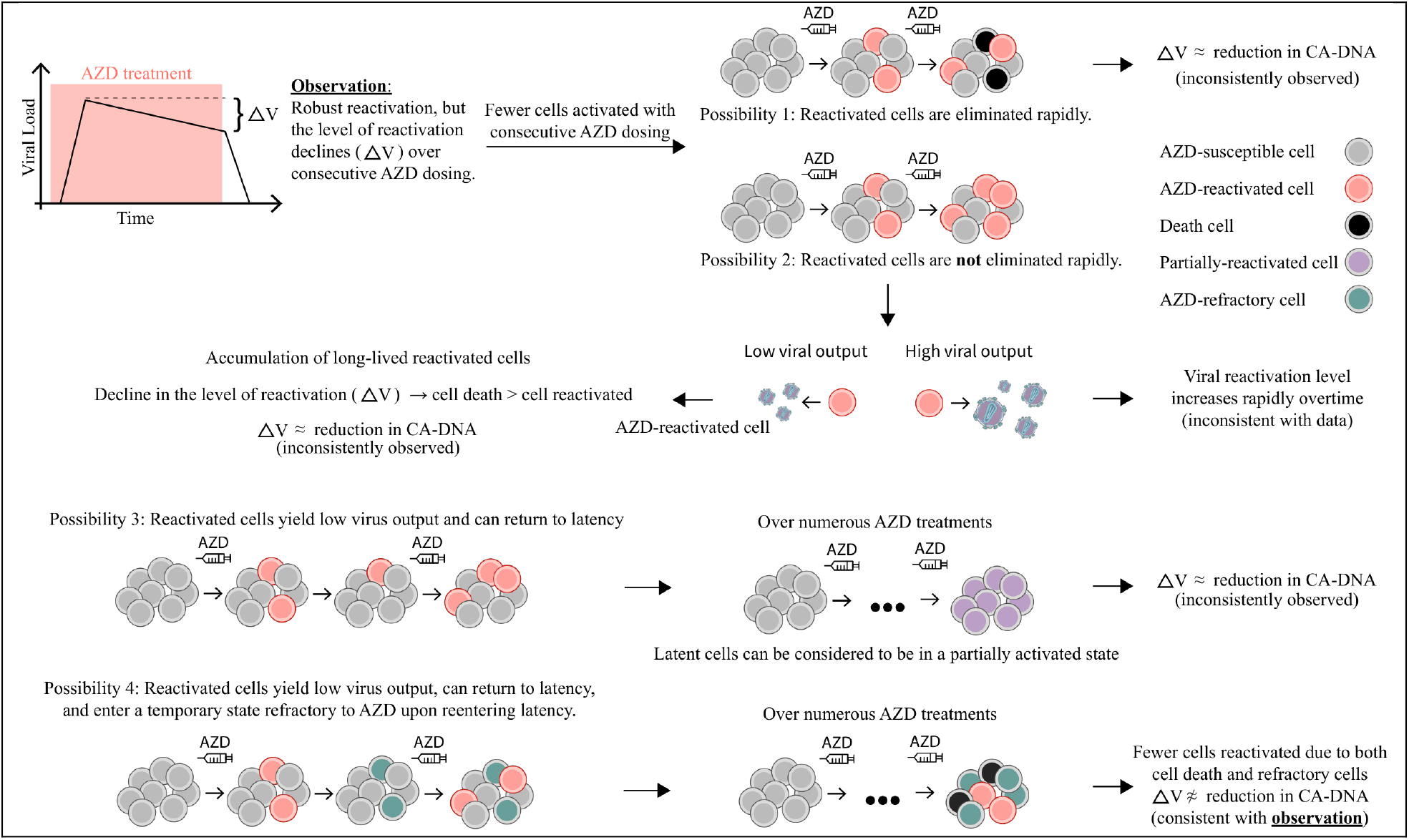
Model components needed to capture the mechanisms of action of AZD. The decline in the level of viral recrudescence (ΔV) and the lack of CA-DNA reduction, together, imply that fewer cells are being reactivated with rapid consecutive doses of AZD. To explain this implication, models require AZD-reactivated cells to differ from productively infected, producing less virus and dying slower, while allowing for latency reestablishment and a temporary refractory state.

## Data Availability Statement

All data and code to reproduce all results will be made available with the publication.

## Ethics Statement

This study involved secondary analysis of longitudinal viral load data from macaques that were previously published. No new animal experiments, sample collection, or interventions were performed for this analysis.

## Author Contributions

Conceptualization: TP, AC, RMR, RK, ASP; Data Curation: MM, AD, AC; Formal Analysis: MM, AD; Funding Acquisition: AC, RMR, RK, ASP; Investigation: TP; Methodology: TP, RMR, RK, ASP; Project Administration: RK, ASP; Resources: MM, AD, AC; Software: TP; Supervision: RMR, RK, ASP; Validation: TP; Visualization: TP; Writing – Original Draft Preparation: TP; Writing – Review & Editing: TP, AC, RMR, RK, ASP.

## Acknowledgements

Portions of this work were done under the auspices of the US Department of Energy under contract 89233218CNA000001 and supported by National Institutes of Health grants P01 AI169615 and R01 OD011095 (to A.S.P.), R01 AI152703 and U54 HL143541 (to R.K.). and Martin Delaney Collaboratory UM1-AI164561 (R.M.R.). T.P. was supported by Director’s postdoctoral fellowship at Los Alamos National Laboratory (20220791PRD2). This research was funded by the National Institute of Allergy and Infectious Diseases and the National Institutes of Health’s Office of the Director, Office of Research Infrastructure Programs, through the following grants: the Emory Consortium for Innovative AIDS Research in Nonhuman Primates (UM1 AI124436 and UM1 AI169662); CARE, a Martin Delaney Collaboratory (UM1 AI126619); the Emory CFAR (P30 AI050409); the Emory National Primate Research Center (P51 OD011132) and R37 AI157862 and R01 AI125064 to A.C. The funders had no role in the study design, data collection and analysis, decision to publish or preparation of the manuscript.

## Supplementary Materials

### S1 Text. Modeling Plasma AZD concentration

We fit a one-compartment PK model to plasma AZD concentration

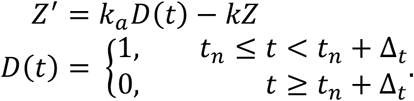

*Z*(*t*) is plasma AZD concentration, *k*_*a*_ is the infusion rate constant over the infusion duration Δ_*t*_, and *k* is the tissue distribution rate constant. The function *D*(*t*) represents the dosing of AZD at time *t*_*n*_. The plasma AZD concentration data, reported in unit of ng/mL, comes from Nixon et al. [30]. However, Sampey et al. reported EC50 for AZD in unit of nM [31], so we convert the unit to nM using AZD molecular weight of 1015.29 g/mol [76]. Model fitting is carried out with MATLAB built-in function *fmincon* using 10 random initial guesses with the function *MultiStart*. Model fit is shown in Fig. S1, and best-fit parameters are presented in Table S1.

**Figure S1.**
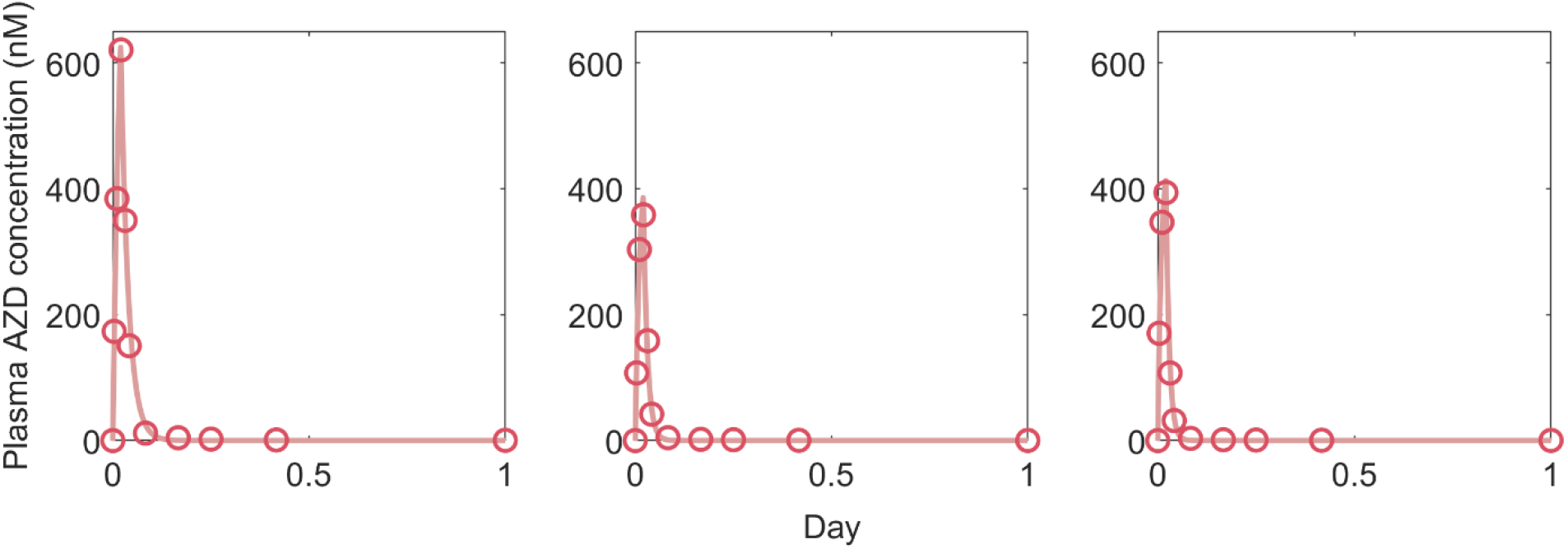
One compartment PK model fit. Red curves are model fit. Open circles are plasma AZD data from three macaques [30].

**Table S1.** Estimated PK parameters for AZD.

| ID | 1 | 2 | 3 |
| --- | --- | --- | --- |
| $k_a$ (day<sup>-1</sup>) | $5.50 \times 10^4$ | $4.17 \times 10^4$ | $6.10 \times 10^4$ |
| $k$ (day<sup>-1</sup>) | 68.92 | 86.81 | 131.21 |
| $\Delta_t$ (day) | 1/48 | 1/48 | 1/48 |

### S2 Text. Mathematical Models M1 – M31

We examined a family of models resulting from the modeling framework introduced by Conway and Perelson [9], which has been shown to accurately capture the HIV and SIV viral load trajectory under different scenarios [11,61–63]. The general model follows Policicchio et al. [61] and Cao et al. [62], where the Conway-Perelson model is extended to include pre-integration infected cells and an exhausted state of effector cells are incorporated (Fig. 2A). Note that the ART regimes in the three groups contain DTG, an integrase inhibitor, necessitating the need for a separation of pre-integrated long-lived and short-lived infected cells [61]. We focused on the period when the SIV-infected macaques were under ART and treated with AZD, so we exclude effector cell exhaustion and details of the infection process (gray boxes in Fig. 2A). The remaining components include productively infected cells *I*, latently infected cells *L*, and effector cells *E*, which make up the basic model (Fig. 2A). The four main variations of the baseline model that we consider are presented in Fig. 2B-E. We present the equations for 31 model iterations (M1 – M31). Parameter definitions and ranges are shown in Table S2. Model fit comparison is presented in Fig. S3 and Table S3.

**Figure S2.**
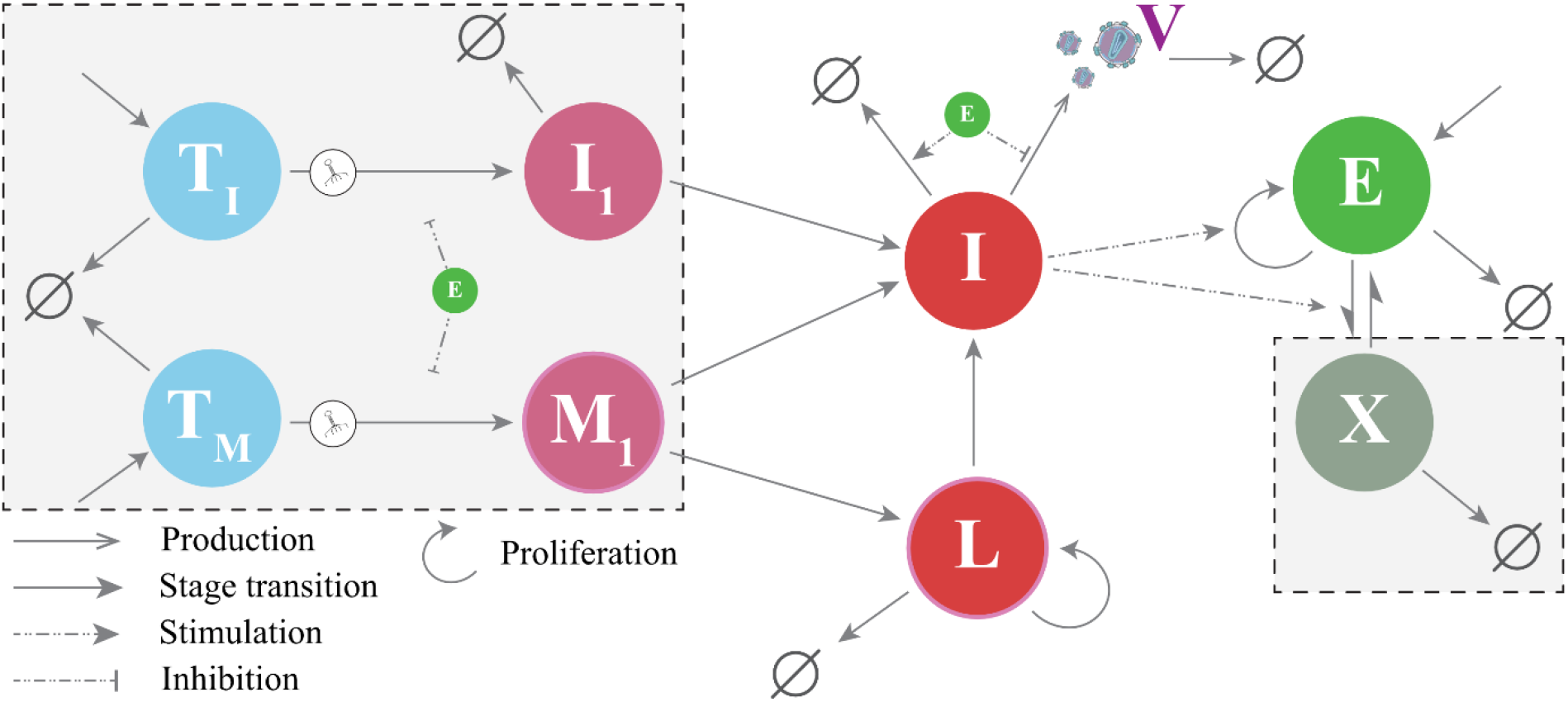
Baseline viral dynamic model. Viruses (V) enter short-(T_I_) and long-lived (T_M_) target cells creating pre-integrated infected short- and long-lived cells I_1_ and M_1_, respectively. After viral integration, these cells become productively infected cells (I), with a fraction of the long-lived cells entering latency (L). Effector cells (E) expand in response to the infection and becomes exhausted (X) over time. During ART, the elements outside of the gray boxes are sufficient to describe viral dynamics.

**M1**.

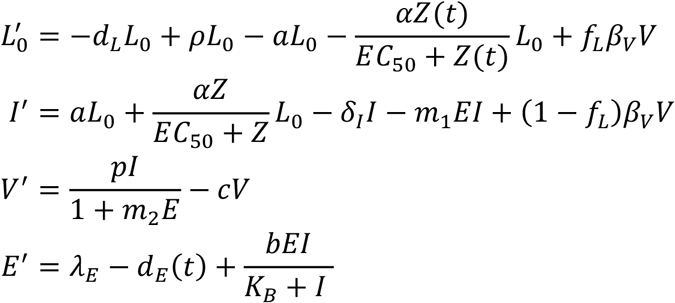

**M2**.

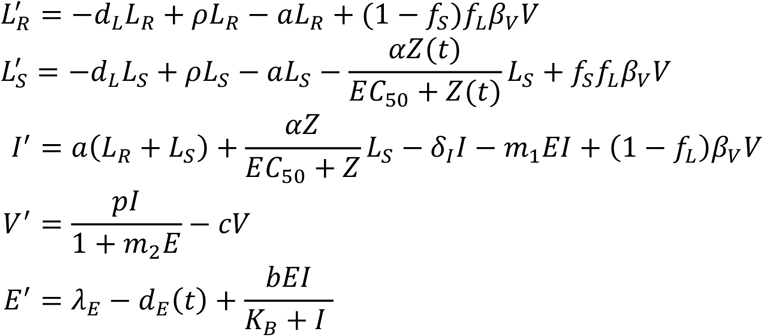

**M3**.

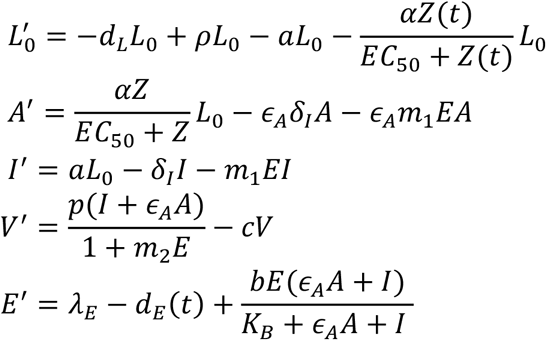

**M4**. We show N=M=3 as the initial demonstration for this variation.

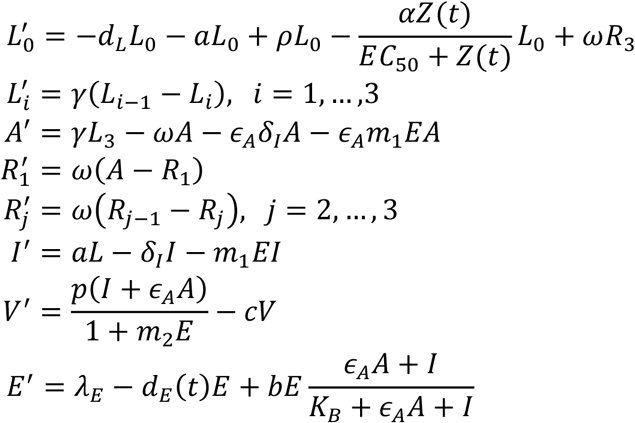

**M5**.

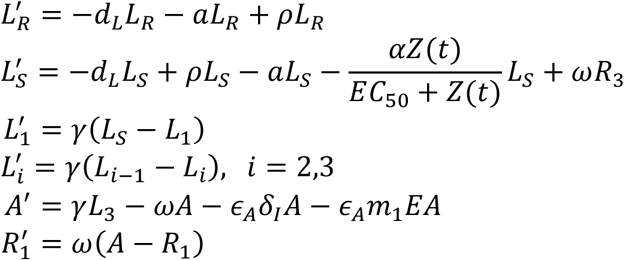

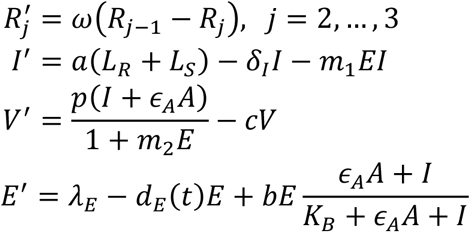

**M6 – M10** as M1 – M5 but without an effector compartment.

**M11 – M12** as M6 – M7 but without active infection (*β*_*V*_ = 0).

**M13 – M15** as M3 – M5 but with active infection.

**M16 – M18** as M13 – M15 but without an effector compartment, or equivalently, as M8 – M10 with active infection.

**M19 – M22** as M9 but with *L*_0_(0) fixed to 10 (M19), 1 (M20), 0.1 (M21) cell per mL, or fitted without random effect (M22).

**M23 – M27** as M22 but with N=M=5 (M23), =7 (M24), =9 (M25), =10 (M26 – lowest BICc), =11 (M27).

**M28** as M26 but without refractory state – AZD-reactivated cells return directly to being susceptible for further induction.

**M29** as M26 but with *ϵ*_*A*_ separated into 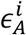, for the effect of infected cell killing and viral production.

**M30** as M10 but with the same modifications as M26.

**M31** as M10 but with the same modifications as M29 (lowest −2LL).

**Figure S3.**
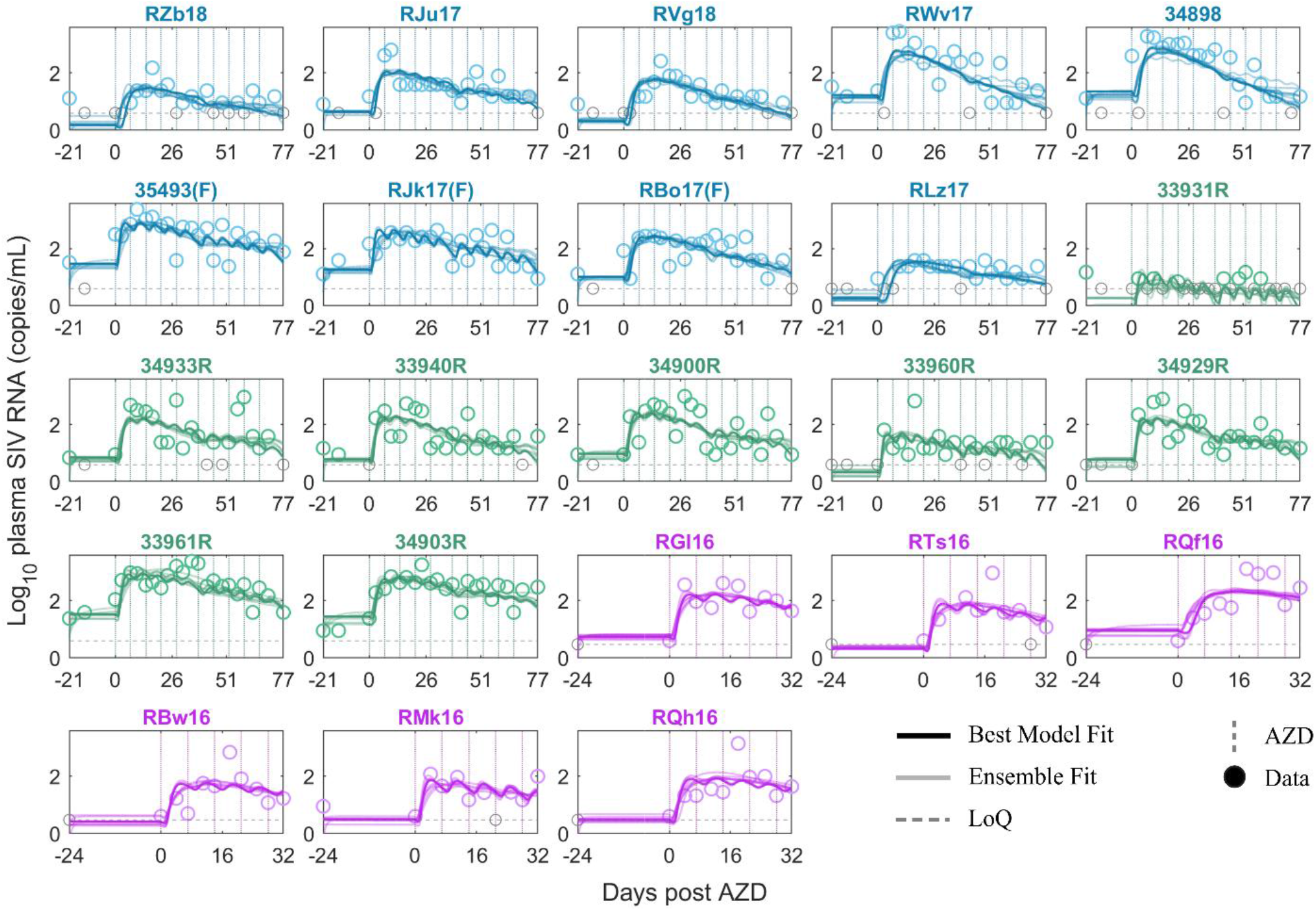
Ensemble and best model fit. Best model (M26) (bolded solid curve). Blue is the group treated with AZD, N-803, and RhmAbs. Green is the group treated with AZD and RhmAbs. Purple is the group treated with anti-CD8α prior to AZD. Ensemble model fit is represented by the transparent curves. Gray circles are data points below the limit of quantification (dotted horizontal gray line). Non-gray circles are data points above the limit of quantification. The dotted vertical lines show AZD administrations.

**Table S2.** Parameter definition. In the Reference column, asterisk indicates fixed values from fitting to data of AZD concentration in blood, and dashed line represents ad hoc ranges. For example, *ϵ*_*A*_ is bounded between 0.01 and 1 to ensure AZD-reactivated cells do not produce more viruses than productively infected cells. The ranges for *m*_2_ and *m*_3_ follow that of *m*_1_. The maximum reactivation rate *α* is assumed to be at least greater than *a*, but no more than 100 per day. Previous estimates [10,103] of the natural reactivation rate <u>without ART</u> are higher than the value used here; however, simulations using M26 with *a* in a similar range (up to 0.1 per day) do not show qualitative differences once AZD begins (except for the steady state value prior to AZD). This reflects that natural reactivation (which is separated from AZD-induced reactivation in our model formulation) is not sufficient to sustain VL during ART. Lastly, note that the estimated parameter values are away from the boundary of these ranges.

| Parameter | Definition & Unit | Range/value | Ref |
| --- | --- | --- | --- |
| $\log_{10} p$ | Viral production rate (day<sup>-1</sup>) | 3 – 5 | [9,11] |
| $L_0(0)$ | Initial size of latent reservoir (cell mL<sup>-1</sup>) | 0.01 – 10 | [96,97] |
| $m_1$ | Effector cell killing rate (day<sup>-1</sup>) | 0.001 – 10 | [9,11] |
| $m_2$ | Effector cell reduction of viral production (cell<sup>-1</sup>) | 0.001 – 10 | - |
| $m_3$ | Effector cell reduction of viral infection (cell<sup>-1</sup>) | 0.001 – 10 | - |
| $b$ | Maximum effector cell expansion rate (day<sup>-1</sup>) | 0.1 – 10 | [9,11] |
| $K_B$ | Density of infected cells needed to reach a half-maximal effector cell expansion rate (mL<sup>-1</sup>) | 0.1 – 10000 | [11,98] |
| $\alpha$ | Maximum reactivation rate of the noncanonical NF-κB pathway by AZD (day<sup>-1</sup>) | 0.01 – 100 | - |
| $\gamma$ | Activation transition rate (day<sup>-1</sup>) | 0.1 – 10 | - |
| $\omega$ | Refractory transition rate (day<sup>-1</sup>) | 0.01 – 10 | - |
| $f_S$ | Fraction of the latent reservoir susceptible to activation by AZD (unitless) | 0.01 – 1 | - |
| $\log_{10} \beta_V$ | Infection rate constant (viral copies<sup>-1</sup> day<sup>-1</sup>) | 3 – 5 | - |
| $\epsilon_A$ | Relative reactivation efficiency ratio (unitless) | 0.01 – 1 | - |
| $\epsilon_A^1$ | $\epsilon_A$ - specific to cytotoxicity (unitless) | 0.01 – 1 | - |
| $\epsilon_A^2$ | $\epsilon_A$ - specific to viral production (unitless) | 0.01 – 1 | - |
| $\epsilon_A^3$ | $\epsilon_A$ - specific to the susceptibility to effector cell cytolytic effect (unitless) | 0.01 – 1 | - |
| $\epsilon_A^4$ | $\epsilon_A$ - specific to stimulation of effector cell expansion (unitless) | 0.01 – 1 | - |
| $\log_{10} \zeta$ | Scaling constant to CA-DNA data (unitless) | 1 – 3 | - |
| $k_a$ | Infusion rate constant (day<sup>-1</sup>) | $5 \times 10^4$ | * |
| $k$ | tissue distribution rate constant (day<sup>-1</sup>) | 90 | * |
| $\Delta_t$ | Infusion duration (day) | 1/48 | * |
| $EC_{50}$ | Concentration of AZD required to reach half-maximal initiation rate (nM) | 7.5 | [31] |
| $a$ | Reactivation rate of latently infected cells (day<sup>-1</sup>) | 0.001 | [9] |
| $d_L$ | Death rate of the latent reservoir (day<sup>-1</sup>) | 0.004 | [64,99] |
| $\rho$ | Latently infected cell proliferation rate (day<sup>-1</sup>) | $a + d_L - \frac{\ln 2}{t_{1/2}}$ | [9,46] |
| $c$ | Viral clearance rate (day<sup>-1</sup>) | 23 | [100] |
| $\delta_I$ | Death rate of productively infected cells (day<sup>-1</sup>) | 1 | [71,101] |
| $\lambda_E$ | Production rate of effector cells (day<sup>-1</sup>) | 1 | [9] |
| $d_E$ | Death rate of effector cells (day<sup>-1</sup>) | 2 | [102] |

**Table S3.** Model fitting comparison order by increasing BICc. Variation refers to the four basic structures in Fig. 2B-D (main text). The “Yes” or “No” indicates the presence (Yes) or absence (No) of an effector compartment, active infection, refractory state, and resistant subpopulation within the model. Shaded rows are the 10 selected models to form the model ensemble. Bolded values are the smallest −2LL and BICc. Note: M23, M24, M25, M27 share the same mechanism as M26, so we omit them from the model ensemble. M19 is the same as M22 and has a lower BICc; however, this is because the fixed value of *L*_0_(0) for M19 happens to be similar to the value estimated for M22. The value of *L*_0_(0) is important, as demonstrated by the difference in BICc across the three models with different fixed values (M19, 20, 21). Thus, we select the model that allows for the possibility of estimating this value for the model ensemble.

| Model | -2LL | BICc | Variation | Effector compartment | Active infection | Refractory State | Resistant subpopulation |
| --- | --- | --- | --- | --- | --- | --- | --- |
| M26 | 851.8 | **933.1** | 3 | No | No | Yes | No |
| M25 | 852.3 | 933.7 | 3 | No | No | Yes | No |
| M27 | 852.5 | 933.8 | 3 | No | No | Yes | No |
| M24 | 853.8 | 935.1 | 3 | No | No | Yes | No |
| M29 | 844.7 | 935.4 | 3 | No | No | Yes | No |
| M23 | 857.1 | 938.5 | 3 | No | No | Yes | No |
| M19 | 864.7 | 939.7 | 3 | No | No | Yes | No |
| M30 | 851.3 | 942.0 | 4 | No | No | Yes | Yes |
| M31 | **842.3** | 942.4 | 4 | No | No | Yes | Yes |
| M22 | 864.1 | 945.4 | 3 | No | No | Yes | No |
| M9 | 865.1 | 949.5 | 3 | No | No | Yes | No |
| M10 | 859.4 | 953.3 | 4 | No | No | Yes | Yes |
| M28 | 878.6 | 959.9 | 3 | No | No | Yes | No |
| M17 | 866.7 | 960.6 | 3 | No | Yes | Yes | No |
| M18 | 861.1 | 964.3 | 4 | No | Yes | Yes | No |
| M16 | 901.6 | 976.6 | 3 | No | No | No | No |
| M8 | 918.2 | 983.9 | 3 | No | No | No | No |
| M7 | 910.7 | 985.8 | 2 | No | Yes | No | Yes |
| M4 | 865.9 | 987.9 | 3 | Yes | No | Yes | No |
| M5 | 862.3 | 993.7 | 4 | Yes | No | Yes | Yes |
| M6 | 929.2 | 994.9 | 1 | No | Yes | No | No |
| M14 | 870.2 | 1011.0 | 3 | Yes | Yes | Yes | No |
| M3 | 910.5 | 1013.7 | 3 | Yes | No | No | No |
| M15 | 867.8 | 1017.9 | 4 | Yes | Yes | Yes | No |
| M13 | 903.2 | 1025.2 | 3 | Yes | No | No | No |
| M2 | 913.2 | 1025.8 | 2 | Yes | Yes | No | Yes |
| M1 | 930.1 | 1033.3 | 1 | Yes | Yes | No | No |
| M20 | 1033.6 | 1108.7 | 3 | No | No | Yes | No |
| M11 | 1102.8 | 1159.1 | 1 | No | No | No | No |
| M12 | 1103.7 | 1169.4 | 2 | No | No | No | No |
| M21 | 1569.8 | 1644.9 | 3 | No | No | Yes | No |

**Table S4.**
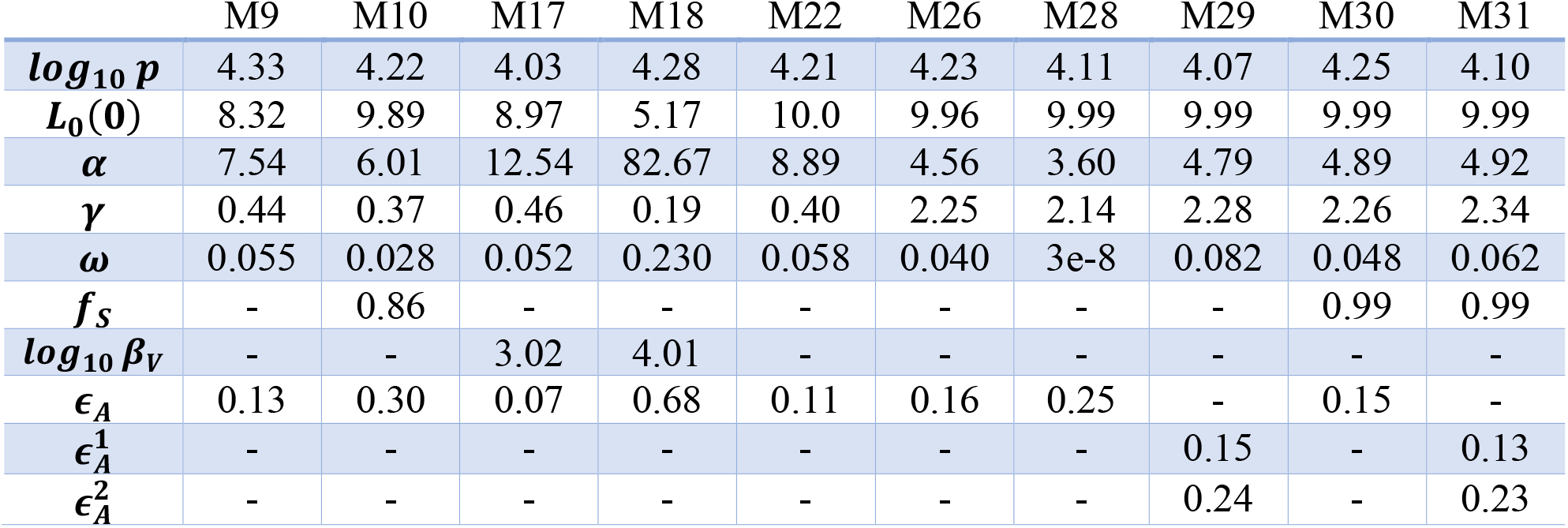
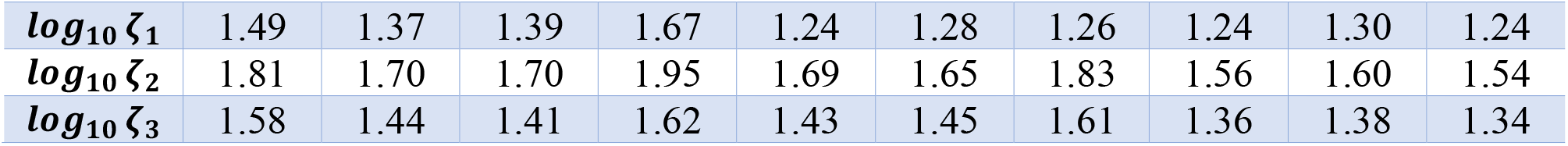
Population parameter estimates of the model ensemble. Dash indicates the model does not contain the parameter. *ξ*_1_ corresponds to the group treated with AZD, N-803, and RhmAbs. *ξ*_2_ corresponds to the group treated with AZD and RhmAbs. *ξ*_3_ corresponds to the group treated with anti-CD8α prior to AZD.

|  | M9 | M10 | M17 | M18 | M22 | M26 | M28 | M29 | M30 | M31 |
| --- | --- | --- | --- | --- | --- | --- | --- | --- | --- | --- |
| **$\log_{10} p$** | 4.33 | 4.22 | 4.03 | 4.28 | 4.21 | 4.23 | 4.11 | 4.07 | 4.25 | 4.10 |
| **$L_0(0)$** | 8.32 | 9.89 | 8.97 | 5.17 | 10.0 | 9.96 | 9.99 | 9.99 | 9.99 | 9.99 |
| **$\alpha$** | 7.54 | 6.01 | 12.54 | 82.67 | 8.89 | 4.56 | 3.60 | 4.79 | 4.89 | 4.92 |
| **$\gamma$** | 0.44 | 0.37 | 0.46 | 0.19 | 0.40 | 2.25 | 2.14 | 2.28 | 2.26 | 2.34 |
| **$\omega$** | 0.055 | 0.028 | 0.052 | 0.230 | 0.058 | 0.040 | 3e-8 | 0.082 | 0.048 | 0.062 |
| **$f_S$** | - | 0.86 | - | - | - | - | - | - | 0.99 | 0.99 |
| **$\log_{10} \beta_V$** | - | - | 3.02 | 4.01 | - | - | - | - | - | - |
| **$\epsilon_A$** | 0.13 | 0.30 | 0.07 | 0.68 | 0.11 | 0.16 | 0.25 | - | 0.15 | - |
| **$\epsilon_A^1$** | - | - | - | - | - | - | - | 0.15 | - | 0.13 |
| **$\epsilon_A^2$** | - | - | - | - | - | - | - | 0.24 | - | 0.23 |
| $\log_{10} \zeta_1$ | 1.49 | 1.37 | 1.39 | 1.67 | 1.24 | 1.28 | 1.26 | 1.24 | 1.30 | 1.24 |
| $\log_{10} \zeta_2$ | 1.81 | 1.70 | 1.70 | 1.95 | 1.69 | 1.65 | 1.83 | 1.56 | 1.60 | 1.54 |
| $\log_{10} \zeta_3$ | 1.58 | 1.44 | 1.41 | 1.62 | 1.43 | 1.45 | 1.61 | 1.36 | 1.38 | 1.34 |

